# PET Brain imaging of α7-nAChR with [^18^F]ASEM: Reproducibility, occupancy, receptor density, and changes in schizophrenia

**DOI:** 10.1101/245118

**Authors:** Dean F. Wong, Hiroto Kuwabara, Andrew G. Horti, Joshua M. Roberts, Ayon Nandi, Nicola Casella, James Brasic, Elise M. Weerts, Kelly Kitzmiller, Jenny A. Phan, Lorena Gapasin, Akira Sawa, Heather Valentine, Gary Wand, Noble George, Michael McDonald, William Kem, Robert Freedman, Albert Gjedde

## Abstract

The α7 nicotinic acetylcholine receptor (nAChR) increasingly has been implicated in normal brain physiology, as well as in neuropsychiatric disorders. The α7-nAChR primarily is located in cerebral cortex and sub-cortical regions, compared to the α4β2 nAChR subtype that has a more subcortical distribution. The highly cortical distribution suggests a role of α7-nAChR in cognition. We expanded the first-in-human PET imaging of α7-nAChR with [^18^F]ASEM from five to 21 healthy non-smoking volunteers and added preliminary evidence of binding in six male patients with schizophrenia. Study aims included 1) confirmation of test-retest reproducibility of [^18^F]ASEM binding in normal volunteers, 2) demonstration of specificity of [^18^F]ASEM binding by competition with DMXB-A, an α7-nAChR partial agonist previously tested in clinical trials of patients with schizophrenia, 3) estimation of [^18^F]ASEM binding potentials and α7-nAChR density in vivo in humans, and 4) α7-nAChR binding in patients with schizophrenia compared to healthy volunteers.

Test-retest PET confirmed reproducibility (>90%) (variability ≤ 7%) of [^18^F]ASEM volume of distribution (*V*_t_) estimates in healthy volunteers. Repeated sessions of PET in five healthy subjects included baseline and effect of inhibition after oral administration of 150 mg DMXB-A. From reduction of binding potentials, we estimated the dose-dependent occupancy of α7-nAChR by DMXB-A at 17-49% for plasma concentrations at 60-200 nM DMXB-A. In agreement with evidence post-mortem, α7-nAChR density (*B*_max_) averaged 0.67-0.82 nM and inhibitor affinity constant (*K*_i_) averaged 170-385 nM. Median *V*_t_ in a feasibility study of six patients with schizophrenia was lower than in healthy volunteers in cingulate cortex, frontal cortex, and hippocampus. Mann-Whitney test identified cingulate cortex and hippocampus as regions with significantly lower median *V*_t_ in patients than in healthy volunteers when a single outlier patient was excluded from analysis (P = 0.02, corrected for multiple comparisons).

## Introduction

As ligand-gated excitatory cation channels, nicotinic acetylcholine receptors (nAChRs) in central nervous system (CNS) are fundamental to physiology (Lukas et al., 1999). The two nicotinic receptor subtypes α4β2 and α7 predominate in CNS among many other nAChR subtypes (Lukas et al., 1999; Gotti and Clementi, 2004). Nicotine binds with high affinity to (α4β2-nAChR (*K*_i_ ~ 1 nM, Anderson and Arneric, 1994) and much lower affinity to α7-nAChR (*K*_i_ ~ 6,000 – 14,000 nM, Davies et al., 1999).

The α7-nAChR subtype is associated with the pathophysiology of specific CNS diseases, including schizophrenia, drug addiction, depression, anxiety, and multiple neurodegenerative disorders (Verbois et al., 2000; Verbois et al., 2002; Woodruff-Pak and Gould, 2002; Albuquerque et al., 2009; D‘Hoedt and Bertrand, 2009; Philip et al., 2010; Hoffmeister et al., 2011; Ishikawa and Hashimoto, 2011; Parri et al., 2011) and therefore is an important target of drug discovery. Multiple post-mortem studies have shown reduced α7 receptors in brain samples from patients with schizophrenia patients, suggesting a role of α7 in schizophrenia (see review by Thomsen et al., 2010). In autoradiography of brains from patients with schizophrenia, reductions of α-[^125^I] BTX binding to α7 were demonstrated in hippocampus, anterior cingulate cortex, and thalamic reticular nucleus (Freedman et al., 1995) but not in orbitofrontal or temporal lobes (Court et al., 1999; Marutle et al., 2001).

The α7-nAChR has emerged as a potential therapeutic target for treatment of schizophrenia (Wallace et al., 2011; AhnAllen, 2012; Geerts, 2012; Wallace and Bertrand, 2013; Young and Geyer, 2013a; Freedman, 2014b; Beinat et al., 2015). In animals, stimulation of α7 is procognitive (Young and Geyer, 2013a; Potasiewicz et al., 2017). In clinical trials with selective α7 agonists, activation of the receptor improved cognitive performance in patients with schizophrenia. Selected drugs (GTS-21, TC-5619, and EVP-6124) show some degree of cognitive improvement in trials of patients with schizophrenia, although none has yet proceeded to late phase or FDA approval. A common trend is the improvement of cognition with an inverted U-shaped dose–response curve, when increasing the dose of α7 agonist above a threshold resulted in loss of drug effect (Wallace and Bertrand, 2015; Lewis et al., 2017). The latest research has revealed a new generation of promising α7-targeting drugs (Beinat et al., 2016).

Despite the importance of the nicotinic receptor system, the physiological and pharmacological roles of specific receptor subtypes in CNS remain poorly understood (Philip et al., 2010; Ishikawa and Hashimoto, 2011; Marrero et al., 2011; Parri et al., 2011). Until recently, the lack of radioligands for quantitative emission tomography imaging of α7-nAChR in humans hampered non-invasive research of this receptor system. Many radiotracers of interest to α7-nAChR (^11^C-CHIBA-1001 (Toyohara et al., 2009), ^11^C-NS14492 (Ettrup et al., 2011; Magnussen et al., 2015, and others) have been tested, but most had suboptimal properties in PET of humans (see Gao et al., 2013 for review). Here, we further tested human subjects with [^18^F]ASEM, the only α7-nAChR PET tracer available to humans with promising imaging characteristics (Gao et al., 2013b; Horti et al., 2014b; Horti, 2015; Coughlin et al., 2017; Hillmer et al., 2017; Wong et al., 2017). Objectives included 1) confirmation of test-retest reproducibility of [^18^F]ASEM binding in brain of normal volunteers, 2) demonstration for the first time of specific binding and target engagement/occupancy in human brain of the partial agonist DMXB-A (GTS-21), 3) estimation for the first time of [^18^F]ASEM binding potentials and α7-nAChR receptor density (*B*_max_) and inhibitor affinity constant (*K*_i_) for human brain in vivo, and 4) a feasibility study of the target role of α7-nAChR in psychosis in six patients with schizophrenia.

## Methods

### Subjects

All studies were carried out with the approval of the Johns Hopkins Institutional Review Board, where all subjects provided written informed consent. Participants were recruited via advertisements within the metropolitan Baltimore region.

Twenty-one healthy, non-smoking control subjects (13 males and 8 females; age range: 18-52 years; means ± SD: 32.6 ± 12.4 years) volunteered for the study. In addition to baseline PET, 12 subjects had second baseline PET (test-retest), and 5 other subjects had a second PET after a single dose of DMXB-A. Only eight of the 12 subjects had successful test-retest PET due to technical problems in either test or retest in four subjects. Of the subjects, five (including two with test-retest from a previous publication by Wong et al. (2014)) were included in this study. Healthy volunteers had no psychotropic treatment history, were free of significant medical or neuropsychiatric disorder, and received no current medications.

To examine the feasibility of measuring a7 nAChR in schizophrenia, six patients diagnosed with schizophrenia (aged 20-31 years; average 25.0 ± 4.3 (SD) years; all males; one smoker only) were recruited. Volunteers with schizophrenia that met DSM-V criteria, with stable antipsychotic medication other than clozapine, were referred by collaborating psychiatrists. Subjects underwent neuropsychiatric characterization including Brief Psychiatric Rating (BPRS) scores. Demographic data for healthy controls and subjects with schizophrenia are listed in Table 1A and 1B. Participants provided information of history of tobacco use, including number of cigarettes per day, and years of tobacco use.

**Table 1A:** Group Demographics

| DIAGNOSIS | CONDITION | N | MALES | FEMALES | AGE<br>(*M/SD*) | AGE<br>(*RANGE*) |
| --- | --- | --- | --- | --- | --- | --- |
| **HEALTHY**<br>**CONTROLS\*** | Baseline | 21 | 13 | 8 | 32.0<br>(12.4) | 18-52 |
|  | Retest | 8 | 6 | 2 | 37.8<br>(11.7) | 20-52 |
|  | Blocking | 5 | 3 | 2 | 22.1<br>(4.5) | 18-28 |
| **PATIENTS WITH**<br>**SCHIZOPHRENIA** | Baseline | 6 | 6 | 0 | 24.8<br>(3.9) | 20-31 |

\*No Healthy controls reported current or past smoking history

**Table 1B:** Patients with Schizophrenia

| Subject ID | Age | Sex | Race | Ethnicity | BPRS –<br>Negative<br>(*M*) | BPRS –<br>Positive<br>(*M*) | Smoker | Cig/day | Medications |
| --- | --- | --- | --- | --- | --- | --- | --- | --- | --- |
| **01** | 27 | M | AA | NH | 2.25 | 1.25 | No | N/A | Aripiprazole |
| **02** | 31 | M | W | NH | 3 | 1 | Yes | 20 | Haloperidol,<br>Benztropine, Sertraline |
| **03** | 22 | M | AA | NH | 2.75 | 2.25 | No | N/A | Risperidone |
| **04** | 25 | M | AA | NH | 2.75 | 1.75 | No | N/A | Quetiapine,<br>Haloperidol, Benztropine |
| **05** | 20 | M | AA | NH | 1.5 | 1.5 | No | N/A | Escitalopram,<br>Benztropine,<br>Aripiprazole,<br>Clonazepam |
| **06 –**<br>**Scan 1** | 24 | M | W | NH | 2.75 | 3.25 | No | N/A | Risperidone, Lorazepam |
| **06-**<br>**Scan 2** | 24 | M | W | NH | 1.0 | 1.0 | No | N/A | Lorazepam |

All subjects (controls and patients with schizophrenia) had negative urinary toxicology tests and provided carbon monoxide (CO) breath samples of < 4 ppm on the imaging day to confirm abstinence for at least 12 hours from tobacco smoking for any subjects (see additional inclusion/exclusion criteria in Supplemental material).

One patient with schizophrenia was tested with PET [^18^F]ASEM while chronically treated with risperidone and then retested with PET without drug for three weeks (discontinued for clinical reasons).

To address possible effects of antipsychotics administered to patients, we determined biodistribution in mice treated for five days with cohorts of placebo vs. risperidone, aripiprazole, or olanzapine, the three most common drugs in the present patient population, at doses equivalent to doses used in humans (Reagan-Shaw et al., 2008; Dawe et al., 2010; McOmish et al., 2012; Ning et al., 2017). Mice were injected IV with [^18^F]ASEM and sacrificed after 60 minutes (detailed Methods and Figure A in Supplemental materials).

### Positron Emission Tomography

All subjects underwent PET with the Siemens/CTI High-Resolution Research Tomography (HRRT: resolution <2 mm). The dynamic imaging continued for 90 minutes with the frame sequences of 4 x 15 sec, 4 x 30 sec, 3 x 60 sec, 2 x 120 sec, 5 x 240 sec, 12 x 300 sec. Each subject received a radial arterial catheter for collection of approximately 40 arterial plasma samples and subsequent HPLC metabolite detection during the 90-minute imaging session. Arterial plasma samples were drawn as rapidly as possible for the first two minutes and then every minute until 10 minutes past injection, every 2 minutes until 20 minutes past injection, and at the times 25, 30, 45, 60, 75, and 90 minutes past injection. Metabolites were measured for the times of 2, 5, 10, 20, 30, 60, and 90 minutes post injection. (See Supplemental data for HPLC methods)

Each subject received 13.8 mCi ±0.23 SEM [^18^F]ASEM synthesized by the method of Gao et al. (2013) with average specific activity of 56,417 Ci/mmol). There were no significant differences in injected radioactivity or specific activity between the controls and patients with schizophrenia. Tracer free fraction in plasma across all subjects averaged 2.79% +/- 0.06 (SD) and was not significantly different in healthy controls and patients with schizophrenia.

Of the healthy volunteers, twelve were retested with [^18^F]ASEM imaging approximately five weeks (mean 41.2 days) after the first PET session, including three reported by Wong et al.(2014). We carefully reviewed the radiometabolite HPLC results and elected to exclude four subjects for technical reasons (e.g., defective capture columns during early development of HPLC procedures). To determine the target engagement (occupancy) by a receptor ligand and to obtain evidence of specific binding of [^18^F]ASEM to α7-nAChR in humans, five healthy subjects additionally underwent a second imaging session after administration of oral 150 mg of the partial agonist DMXB-A (also known as GTS21), forty minutes before the PET tracer injection. Radial arterial samples for detection of concentration DMXB-A were obtained during PET imaging at 70, 100, and 130 minutes post-dose. Concentrations of DMXB-A in subjects’ plasma samples were determined quantitatively by HPLC with a modification of the previously reported method (Kem et al., 2004). (Details in supplemental information).

Menstrual phase for the women was determined by a progesterone level using levels exceeding 2 ng/ml, indicating luteal phase. For two women (one in luteal phase and one in follicular phase), menstrual phase was determined by self-reported menstrual history, as progesterone samples were not available.

## PET Analysis

### Volumes of interest

Volumes of interest (VOI) were selected automatically in individual subject SPGR MRI volumes using FSL (FMRIB Software Library; Jenkinson et al., 2012) FIRST tool (Patenaude et al., 2011) for subcortical regions and Freesurfer tool (Fischl et al., 2004) for cortical regions. Automated VOI were manually edited to suit the structures of interest, using a locally developed VOI tool (VOILand). Refined VOI were transferred from MRI to PET spaces according to MRI to PET coregistration parameters obtained from the co-registration module of SPM12 (Statistical Parametric Mapping; Ashburner and Friston, 2003). The VOI of PET space were applied to PET frames to obtain time-activity curves (TAC) of regions. Head-motion correction (HMC) was performed with the image co-registration module of SPM12.

Total volumes of distribution (*V*_t_) and test-retest variability were estimated for all 21 cortical VOI (Figure 1A and B). In addition, the full VOI set was used in the calculation of dose-dependent inhibition (Figure 2), as well as for the averaged estimates of maximum binding capacity (*B*_max_; Figure 3), and inhibitor affinity constant (*K*_i_; Figure 4). For analysis of *V*_t_ differences (in patients vs. control subjects, Figure 5) and potential correlation with BPRS score (see Supplementary Material) we chose three cortical regions implicated in autoradiography with [^125^I]α-bungarotoxin ([^125^I]BTX) of brain of patients with schizophrenia postmortem, previously reported with significant reductions in hippocampus (~70%, Freedman et al., 1995), frontal cortex (40%, Guan et al., 1999), and cingulate cortex (54%, Marutle et al., 2001).

**Fig. 1:**
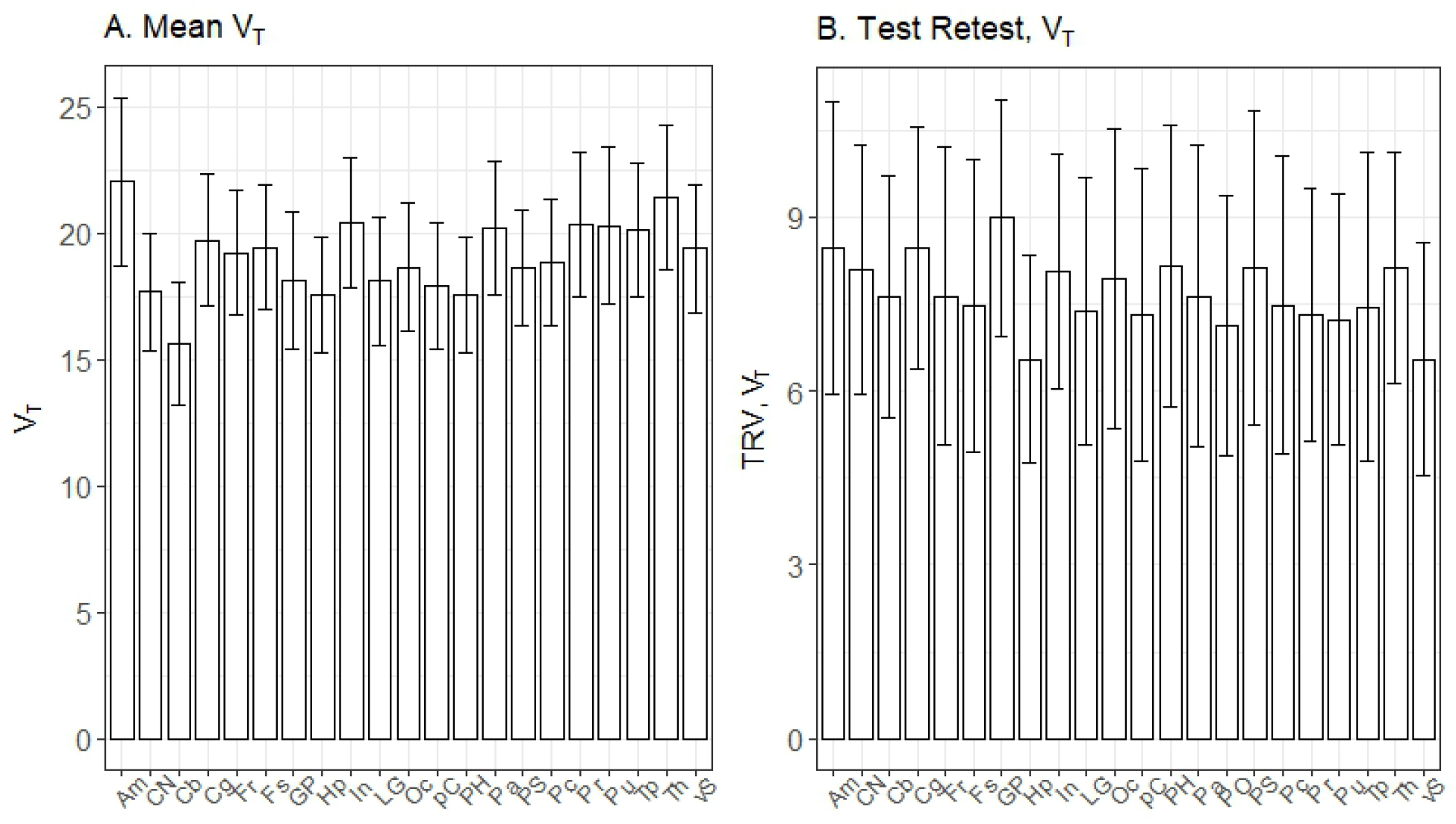
Estimates of *V_t_* and Test-Retest Variability for *V_t_*. **1A:** Regional estimates of *V_t_*, for all healthy control subjects (n=21). **1B:** Test-Retest variability of *V_t_* in eight healthy controls. For both panels, bars show mean +/- SD. **Regional Abbreviations:** Am: amygdala; CN: caudate nucleus; Cb: cerebellum; Cg: cingulate; Fr: frontal lobe; Fs: fusiform gyrus; GP: globus pallidus; Hp: hippocampus; In: insula; LG: lingular gyrus; Oc occipital lobe; pC: paracentral; PH: Parahippocampus; Pa: parietal lobe; PS: postcentral gyrus; Pc: precentral gyrus; Pr: precuneus; Pu: putamen; Tp: temporal lobe; Th: thalamus; vS: ventral striatum.

**Fig. 2.**
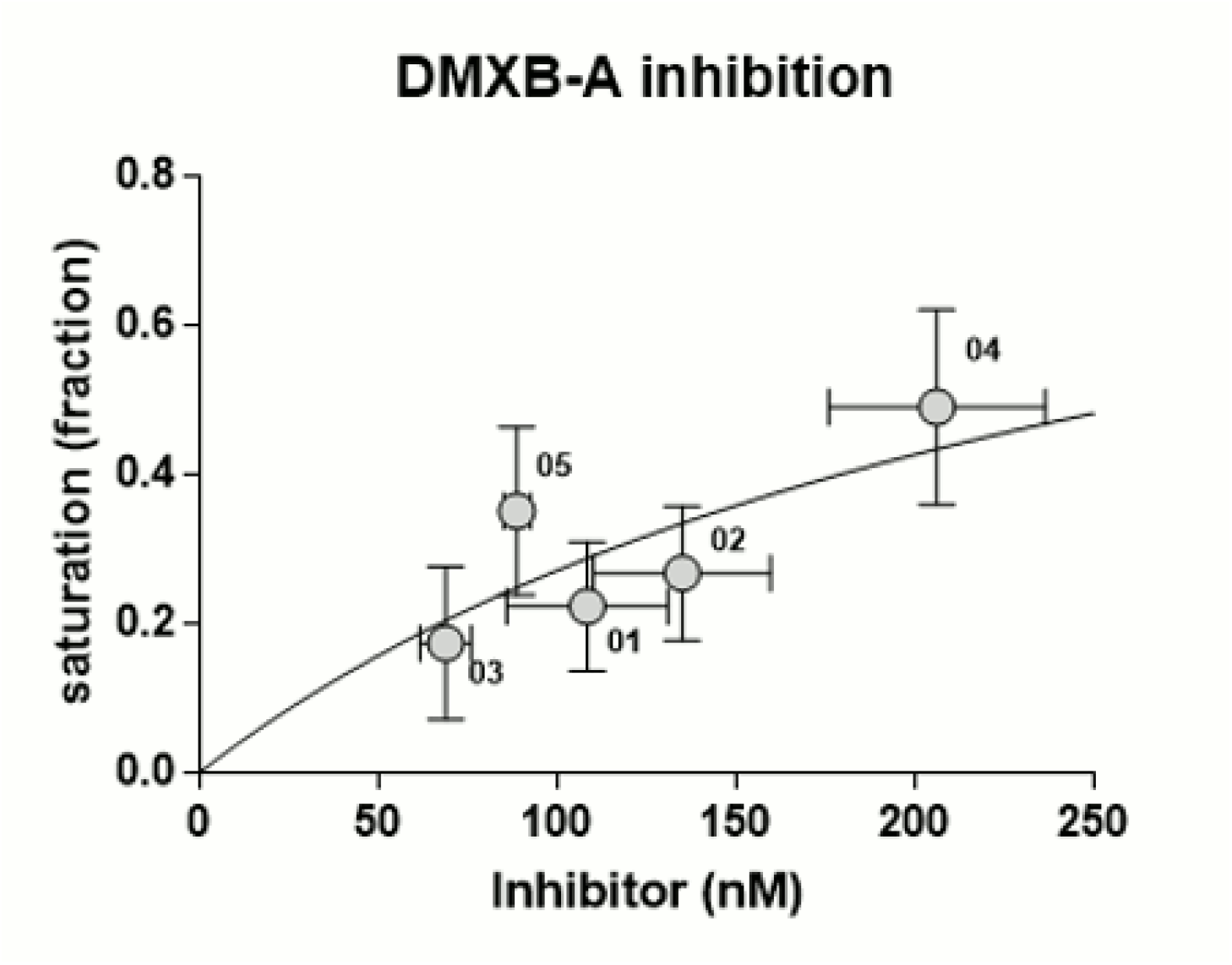
Dose-Dependent Inhibition with DMXB-A level. The saturation, *s*, reflects the fraction of receptor occupied by the blocking agent, DMXB-A. Each point on the curve represents individual measurements of the plasma level of DMXB-A, determined by mass spectroscopy, and the fraction of receptors occupied by DMXB-A, estimated by the Extended Inhibition Plot. The horizontal error bars represent the S.E.M from three measurements, and the vertical error bars reflect the S.E.M from the linear analysis of the Extended Inhibition plot. These data were fit to the Michaelis-Menten equation, with *O_max_* assigned to a value of 1; (i.e, 100% occupancy, or a saturation fraction, *s* = 1). This is typical for receptor occupancy studies with PET. However, further studies at higher plasma concentrations may indicate a lower maximal occupancy, due to the unique nature of the α7 subunit, as stated in the Discussion.

**Fig. 3.**
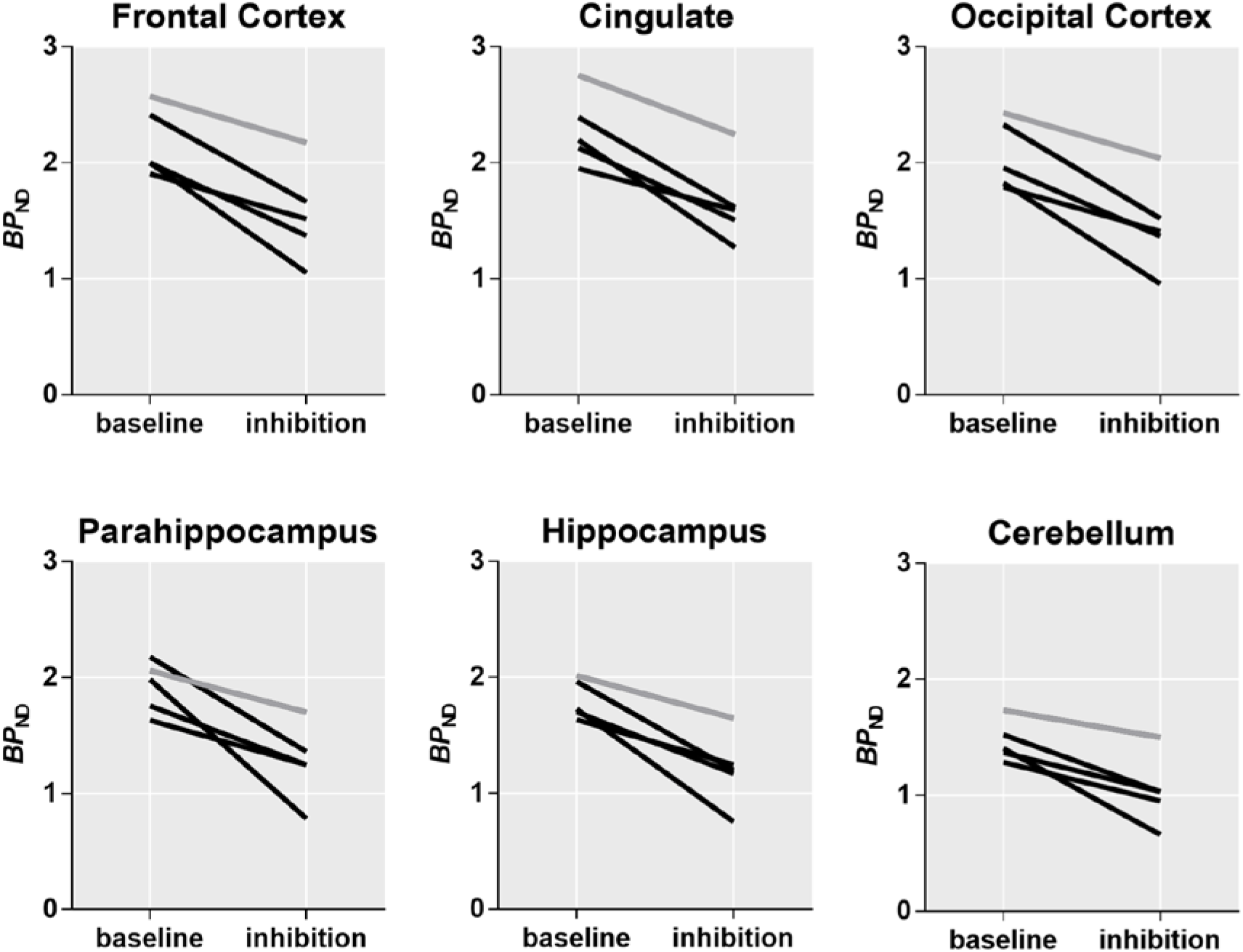
Regional Binding Potentials (*BP_nd_*) The analysis using the Extended Inhibition plot revealed that the binding potential, *BP_nd_*, decreased significantly in response to competition with DMXB-A by 31% in average when compared to baseline. Each line on the graphs of six representative regions of the original 21 regions displays the estimates of BPND in individual subjects at baseline and inhibition condition.

**Fig. 4.**
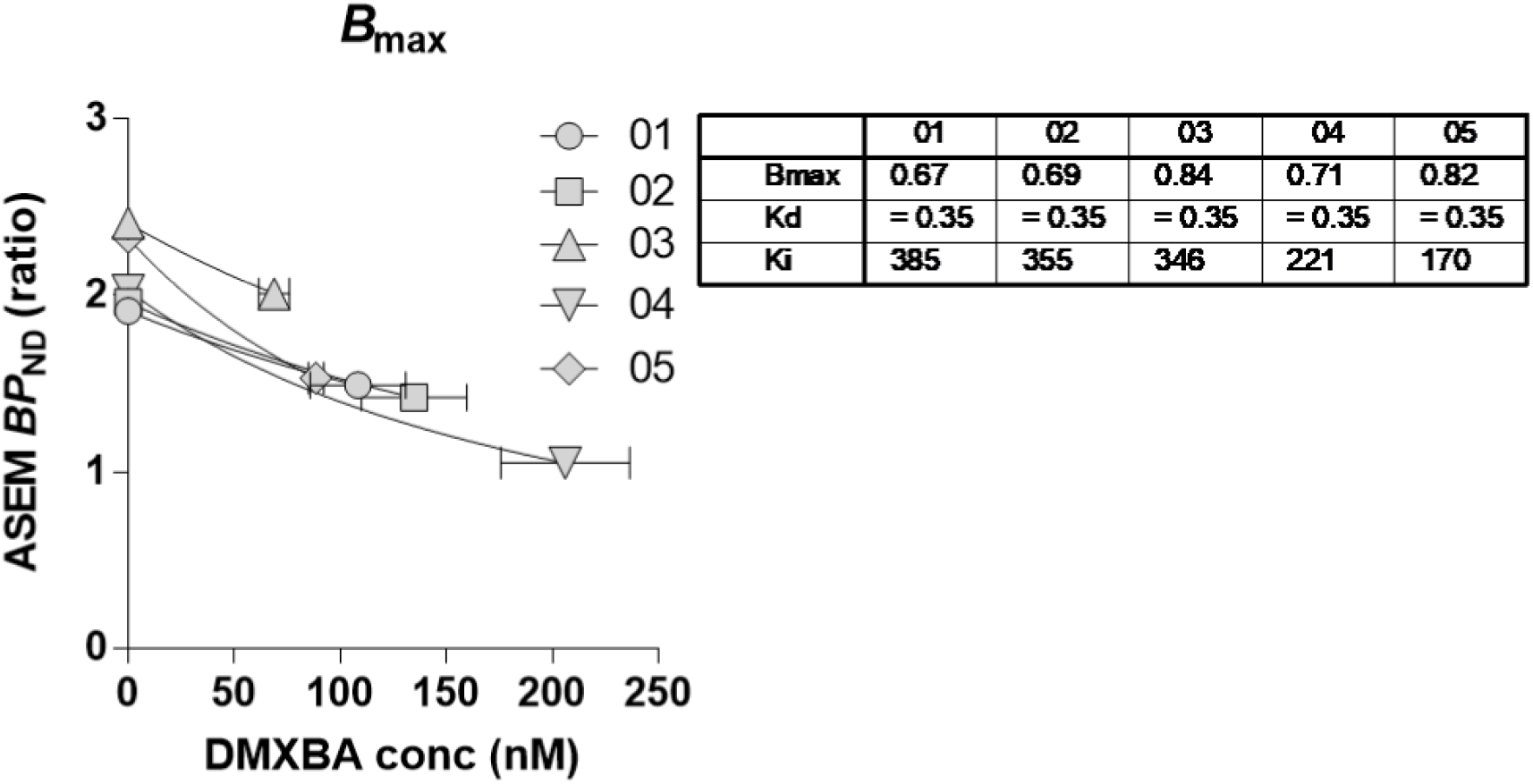
*B_max_* estimates. Average *B_max_* across all regions (21 VOIs total) were computed from the known *BP_nd_* estimates, the plasma concentration of DMXB-A and the receptor affinity, *K*_d_, which was calculated as 0.35 nM based on known values from ^125^I α-bungarotoxin *in vitro* studies. The analysis reveals that the average *B*_*max*_ across all regions ranged from 0.76-0.96 nM in the five subjects, and the *K_i_* ranged from 170-385 nM.

**Fig. 5.**
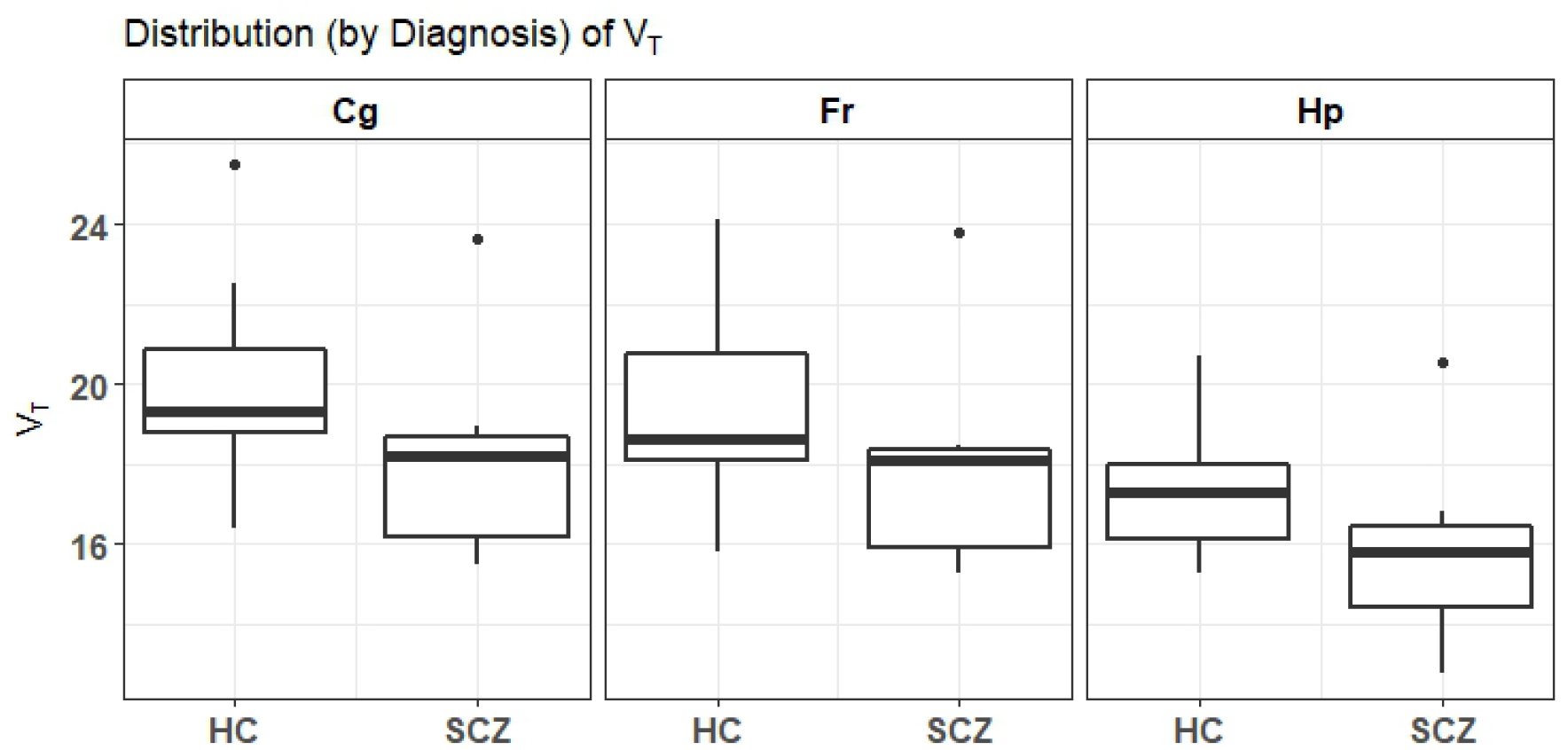
Distribution of *V*_t_ in healthy patients and controls. Boxplots of the distribution of *V*_t_ in healthy controls (HC) and patients with schizophrenia (SCZ), in Cingulate Cortex (Cg), Hippocampus (Hp), and Frontal Cortex (Fr). The single points above the limits of the boxplot are outliers.

### Kinetic analysis

Quantification of [^18^F]ASEM steady-state volumes of distribution (*V*_t_) proceeded as previously published (Wong et al., 2014). Specifically, the plasma reference graphical analysis (PRGA) was used to obtain regional *V*_t_ values that served as primary outcome variable for test-retests and analyses with age, sex and neuropsychiatric testing, as well as serving as secondary outcome variable for estimates of the reference volume of non-displaceable bound radioligand (*V*_nd_), competitor saturation of receptors (*s*), binding potentials (*BP*_nd_), and receptor density (*B*_max_). We completed receptor density (*B*_max_) and affinity (IC_50_, *K*_i_) estimates of α7-nAChR in three steps in five subjects by means of DMXB-A inhibition.

We estimated the occupancy of the partial agonist inhibitor by means of estimates of the radioligand’s volume of non-displaceable distributed tracer in the baseline (*V*_nd_), obtained from baboon brain in vivo (Horti et al., 2014), using the conventional Inhibition Plot (IP) of Gjedde and Wong (2000). We compared the value of *V*_nd_ obtained from the decline of the magnitude of the total volume of *V*_t_ in in the presence of two different concentrations of SR180711, of which we used the value associated with the lower dose to avoid affecting the value of *V*_nd_ by changes of protein binding in brain tissue or circulation or both (see Supplemental Figure B illustrating the *V*_nd_ determination in baboon brain).

Applied to regional volumes of distribution (*V*_t_), the Inhibition Plot yielded reference volumes (*V*_nd_) and degrees of saturation (*s*), according to the equation (Phan et al., 2017),

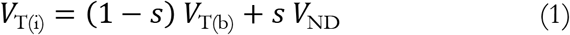

where *V*_t (b)_ symbolizes the volume of distribution at baseline, *V*_T (i)_ the volume at inhibition in presence of DMXB-A which competes with the radiotracer for binding to the binding site. According to Equation (1), when the regional estimates of *V*_t_ in the inhibition condition were plotted as functions of the estimates in the baseline condition, the solution yielded the reference volume, *V*_nd_, and the fraction of receptors occupied by the competitive drug, *s*.

The initial analysis revealed *V*_t_ estimates before and after DMXB-A inhibition of similar magnitudes, compensating for values of *V*_nd_ that may be different in the inhibition and baseline conditions. To resolve this issue, we applied the Extended Inhibition Plot equation (EIP; (Phan et al., 2017)) that relates the radioligand distribution volumes at inhibition and baseline as follows,

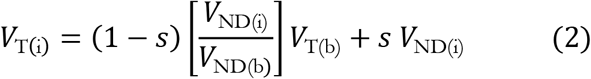

where values of *V*_nd_ at baseline and inhibition are different, denoted *V*_nd (b)_ and *V*_nd (b)_ at baseline and inhibition, respectively. The Extended Inhibition Plot (EIP) yielded the value of *V*_nd_ in the inhibition condition (*V*_nd(i)_), when the true reference volume at baseline (*V*_nd(b)_) is known, such that estimates yielded by EIP depend on a measure of the magnitude of the reference volume.

Volumes of distribution (*V*_t_ and *V*_nd_) yielded binding potentials (*BP*_nd_). We estimated the binding potentials of five subjects with DMXB-A inhibition in the baseline and at inhibition, using the values of *V*_t(b)_ and *V*_nd(b)_ in the baseline condition, and the values of *V*_t(i)_ and *V*_nd(i)_ at inhibition obtained by means of equation (2), in the unblocked and DMXB-A inhibited states, with the equations,

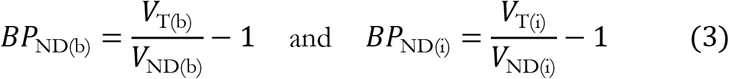

where the *BP*_nd (b)_ and *BP*_nd(i)_ terms symbolize the binding potentials at baseline and inhibition, respectively.

The binding potentials (*BP*_nd_) and inhibitor concentrations (*C*_1_) yielded the *B*_max_ and *K*_i_ (IC_50_) estimates of the receptor and inhibitor, respectively. We completed receptor density (*B*_max_) and affinity of the competitor (IC_50_, *K*_i_) calculation of α7-nAChR in five subjects by means of DMXB-A inhibition, using the values of *BP*_nd_ of baseline and inhibition that led to the novel (never previously published) in vivo estimate of the magnitude of the receptor density (*B*_max_) and affinity (*K*_i_) of nAChR in humans, using the equation,

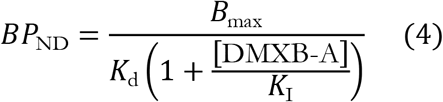

where the receptor density is determined relative to the affinity of the radioligand to the receptor, *K*_d_, with known values of the inhibitor concentration [DMXB-A] and the receptor’s affinity for inhibitor, *K*_I_. For this calculation we used a calculated value of ASEM *K*_d_ of 0.35 nM (established from prior measure of *K*_I_ of 0.4 nM for [^123^I] bungarotoxin).

### Schizophrenia

As a feasibility study, we compared six patients with schizophrenia to a subset of 15 healthy volunteers matched for age, within three specific brain regions: hippocampus and cingulate and frontal cortices. In this subset the mean age of healthy volunteers was 23 ± 4 (SD) years, range 18-30; and patients with schizophrenia mean age was 24 ± 4 (SD) years, range 20-31 years.

## Results

### Regional Volumes of Distribution

Regional estimates of volumes of distribution (*V*_t_) in selected regions are shown in Figure 1A. We tested the test-retest variability of the *V*_t_ estimates in eight healthy subjects, six males (average age 41) and two females (average age 24). Across all cortical and subcortical VOIs, the average test-retest variability was 7.8% ± 0.6% (SD, see Figure 1B), ranging from 6.5% to 8.9% (repeatability of > 90%).

### Receptor Kinetics

The reanalysis of inhibition of [^18^F]ASEM binding in baboon (Horti et al., 2014) yielded the *V*_t_ of non-displaceable tracer accumulation at baseline (*V*_nd (b)_), In the baboon, the *V*_t_ of [^18^F]ASEM declined markedly in response to blocking with the high affinity inhibitor SSR180711. The reference volume *V*_nd_ assessed by means of the standard Inhibition Plot (IP) is presented in Figure 2, with the estimate of *V*_nd (b)_ of 5.4 ml/cm^3^ at the lower dose of the SR180711. We note that the magnitude of *V*_nd_ is a property of the tracer [^18^F]ASEM that we assume to be representative of mammalian brains in general.

We used the value of *V*_nd_ of 5.4 ml/cm^3^ as the value of *V*_nd (b)_ in the Extended Inhibition Plot (EIP), required when *V*_t_ values failed to yield evidence of inhibition of radioligand binding by DMXB-A in human brain. In five subjects, the value of (*V*_nd(b)_ yielded estimates of saturation (*s*) and inhibition reference volume (*V*_nd(i)_), as shown in Figure 2 of the individual estimates of *s* as function of the concentration of DMXB-A in arterial plasma, confirming the dose-dependent inhibition exerted by DMXB-A. Estimates of regional binding potentials (*BP*_nd_) declined significantly in 6 of the 21 representative regions after blocking with DMXB-A, as shown in Figure 3. The average values for all 21 regions of receptor density (*B*_max_) and average values of the receptor’s affinity (*K*_i_) to DMXB-A for ranged from 0.67-0.82 nM for *B*_max_ and from 170 to 385 nM for *K*_i_, as presented in Figure 4. Regarding the choice of the *V*_nd (b)_ the standard error of the *V*_nd (b)_ estimate in baboon of 2.7 ml/cm^3^ corresponded to ranges of human *B*_max_ estimates of 0.4-1.9 nM and DMXB-A KI estimates of 138-363 nM.

### Menstrual Cycle

The menstrual cycle phase (luteal vs. follicular) was determined for the eight healthy control females. Estimates of *V*_t_ did not differ significantly in relation to cycle phase. The lower ages of women volunteers precluded analysis of different changes of *V*_t_ with age in men and women.

### Schizophrenia

To evaluate the feasibility of studying patients with schizophrenia, we compared patients with healthy control subjects. In the three preselected regions (cingulate and frontal cortices, hippocampus) examined for group differences between patients and the age-matched control subjects, the median *V*_t_ values were lower in patients than in age-matched healthy volunteers (Figure 5). Potential outliers, one a control subject and one a patient, were identified (outlier points marked in boxplots of Figure 5). Using the Mann-Whitney U-test, we tested for significant differences of median *V*_t_ values between control subjects and patients, before and after excluding outliers. The results, with corrections for multiple comparisons, are given in Table 2. In the cingulate cortex and hippocampus, the difference of *V*_t_ values was significant after exclusion of outliers and correction for multiple comparisons (P = 0.02 in both regions).

**Table 2:**
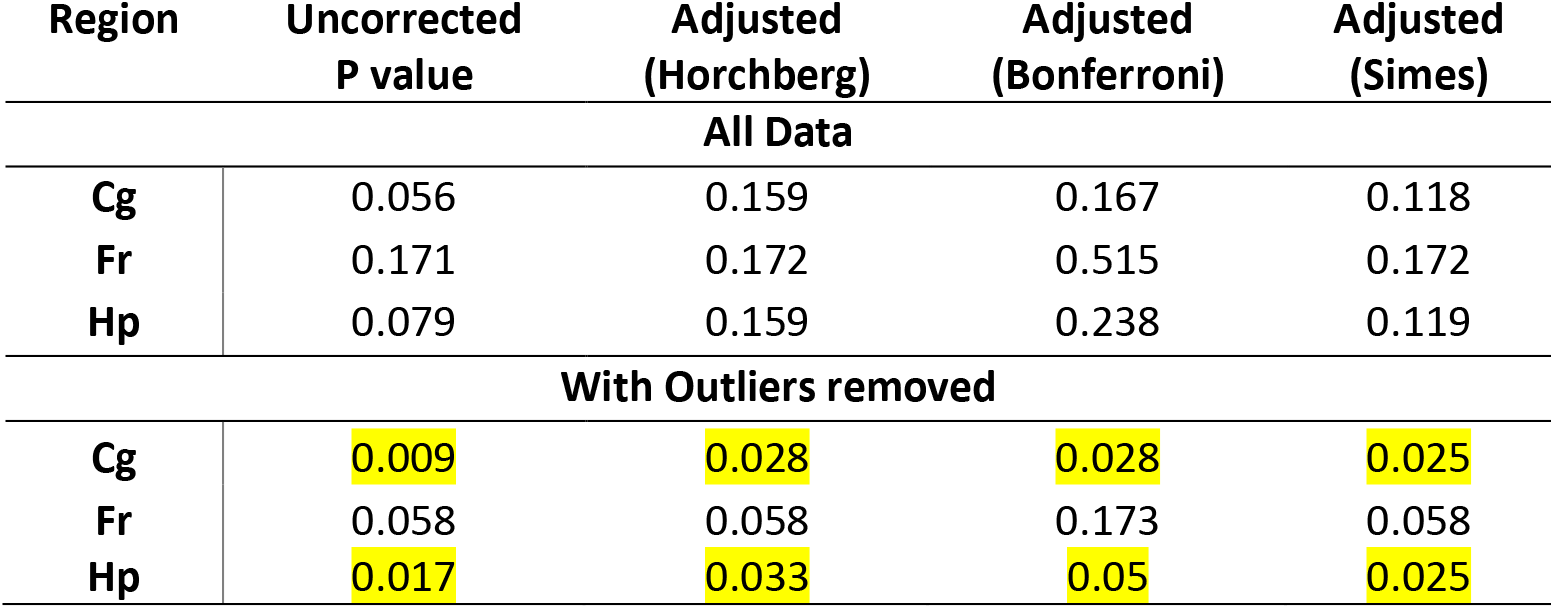
Corrected and uncorrected P values from *V_t_* in controls vs. patients. Results (uncorrected P values and corrected by different methods) of Mann-Whitney tests for *V*_t_ in Healthy Controls vs. Patients with Schizophrenia. Adjusted p values were obtained by the STATA package *qqvalue* using three different methods of adjustment (Newsom, 2010; specific methods are Horcberg, 1998, standard Bonferroni correction and Simes, 1986).

The correlation of estimates of *V*_t_ with scores of BPRS did not reach statistical significance although there were positive trends for values in cingulate and frontal cortices in five subjects after outlier analyses (see Supplementary Figure C for details).

### Antipsychotic medication

The biodistribution in mice revealed no significant effects on [^18^F]ASEM binding of aripriprizole, risperidone, or olanzapine (see Supplemental materials). As described in methods, the patient with schizophrenia chronically treated with risperidone had repeated PET after a 3-week drug-free period, imposed for clinical reasons. The *V*_t_ estimates did not differ by more than 7% (i.e., within the test-retest variability of control subjects (Figure 1B)) in any brain region for the on- and off–risperidone comparison. (Supplementary Figure C, points labelled “on” and “off’).

## Discussion

### Subjects

We extended the first-in-human report of (Wong et al., 2014) from five to 21 healthy subjects (eight test-retest sessions, five sessions with inhibition by DMXB-A of [^18^F]ASEM binding, and six baseline sessions) and added a preliminary study of six patients with schizophrenia, with one patient receiving test-retest PET in the absence and presence of chronic risperidone medication.

The results demonstrate excellent test-retest repeatability of [^18^F]ASEM PET of more than 90%. In the population of 21 healthy non-smoking volunteers, when we plotted tracer distribution volume *V*_t_ against age we observed an increase with of *V*_t_ with age. While our smaller sample and lack of a wide age distribution limited our ability to test this relationship formally with regression analysis, the trends in our data match the findings from a larger age range in another set of subjects (Coughlin et al., 2017; which had only 2 subjects in common which we provided to the Coughlin paper previously). The preliminary findings of values of *V*_t_ vs age are shown in Supplementary Figure D.

We were unable to determine sex differences because of the different age ranges. Of the eight women, none showed a specific effect of menstrual cycle on estimates of *V*_t_.

### Receptor binding potentials

Estimates of receptor density (*B*_max_) are based on determination of binding potentials *BP*_nd_ (Gjedde et al., 2005). Importantly, for the first time, we present evidence that competitor occupancy, apparent inhibitor affinity (*K*_i_), and receptor density (*B*_max_) of human α7-nAChR can be measured, based on the assumptions presented. We determined regional estimates of [^18^F] ASEM binding as the first step towards complete receptor quantification. In addition to regional values of *V*_t_, the receptor quantification required knowledge of the magnitude of *V*_t_ of the tracer in a region of brain tissue with no specific binding. Previously, no such reference volume has been identified for this tracer with prevalent binding in cerebral cortex. As the first PET study with blockade of α7-nAChR in human brain in vivo, this study took advantage of the original development of the well-documented partial agonist DMXB-A at the University of Florida (Kem, 2000; Kem et al., 2004) and the subsequent successful testing in patients with schizophrenia at the University of Colorado (Olincy et al., 2006; Freedman, 2014).

### Receptor occupancy

Recent studies of mouse and baboon brains revealed significant blockade of the α7-nAChR by the high affinity partial α7-nAChR agonists SSR180711 and DMXB-A in vivo (Horti et al., 2014a; Wong et al., 2014), but effective inhibition or competition has not yet been confirmed in humans. As a candidate drug, DMXB-A was tested in clinical trials of patients with schizophrenia, Alzheimer’s disease, or ADHD (Freedman et al., 2008; Freedman, 2014), and the drug was used here to test target engagement/occupancy in vivo at the α7-nAChR and to demonstrate specificity of [^18^F]ASEM binding to these receptors.

Previously, we demonstrated dose-dependent tracer inhibition in rodents by DMXB-A with cerebellum as reference region (appropriate in rodents but not human brain) (Wong et al., 2014). Similarly, in the present study, blockade with the highest dose of DMXB-A allowed in humans (150 mg orally) exhibited competition with binding of [^18^F]ASEM. The occupancy of receptors by DMXB-A displayed a relatively monotonic relationship with plasma levels of DMXB-A.

We used the Extended Inhibition Plot (EIP) under the assumption that the apparently absent inhibition of receptor binding would be due to factors such as change of free fractions of the radioligand in circulation or brain tissue or both, with the consequence that the value of *V*_nd_ would differ between the baseline (*V*_nd (b)_) and inhibited (*V*_nd (a)_) states, as reported by Phan et al. (2017). Using EIP and the value of *V*_nd(b)_ of 5.4 ml/cm^3^ obtained from moderate blockade of the α7-nAChR in baboon brain, reported by Horti et al. (2014) as a simple reduction in slope and reanalyzed by the standard Inhibition Plot (IP) here, we obtained DMXB-A occupancies in the human subjects in the range of 17-49%. Future studies with new α7-nAChR drugs will further test the occupancy and the accuracy of the resulting estimates of receptor density and affinity in human brain in vivo.

With the assumptions and translation from autopsy protein and wet tissue based receptor densities (*B*_max_) of α7-nAChR, we find remarkable agreement among the *B*_max_ estimates that we regard as another innovative aspect of the present study. As 150 mg is the highest single oral dose allowed under the DMXB-A IND based on historical information, we elected not to test lower doses, because of the modest occupancy, compared to occupancy of drugs such as antipsychotics, and more recently glycine transporter (GlyT1) ligands (Martin-Facklam et al., 2013; Wong et al., 2013), the dose dependent occupancy with DMXB-A is evidence of specificity of the radiotracer [^18^F]ASEM in human brain. The occupancy is further evidence of the potential usefulness of [^18^F]ASEM for therapeutic occupancy trials, as these are somewhat complicated due to the inverted-U shaped dose effect of α7-nAChR partial agonists (Young and Geyer, 2013).

The α7-nAChR receptor has five binding sites among the five α7 subunits and is maximally activated when at least two binding sites are occupied by an agonist. At higher doses, agonists bind at up to all five binding sites and progressively desensitize the α7. Moreover, α7 partial agonist drugs such as DMXB-A and others produce their clinical effects at lower concentrations than would be expected from *in vitro* functional experiments where peak currents are recorded (Prickaerts et al., 2012; Papke, 2014; Preskorn et al., 2014). The co-agonistic mechanism of *in vivo* action of α7 partial agonist suggests that DMXB-A, at the clinically effective dose, binds at one of five binding sites of α7, while brisk activation of the receptor will, however, occur upon exposure to endogenous ACh. Thus, *in vivo*, a single molecule of DMXB-A plus one molecule of ACh are sufficient to trigger the opening of the α7-nAChR channel (Williams et al., 2011; Prickaerts et al., 2012; Papke, 2014; Preskorn et al., 2014). Therefore, in the PET occupancy studies with the DMXB-A-treated humans, the binding of [^18^F]ASEM is expected to be inhibited by as little as 20-40% (1/5^th^ - 2/5^th^) instead of 100%. This co-agonistic mechanism agrees reasonably with our PET [^18^F]ASEM <u>inhibition by</u> DMXB-A in humans at 17-49% with a maximum clinical dose of DMXB-A (150 mg). Full blocking of [^18^F]ASEM with α7 agonists occurs at higher doses, as observed in mice with 20 mg/kg DMXB-A (Horti et al., 2014a; Wong et al., 2014) and in baboons administered 5 mg/kg of the more highly potent SSR180711 that is unavailable in the US (Horti et al., 2014).

### Receptor Density and Tracer Affinity

The use of the EIP with the DMXB-A inhibition and the use of the estimate of *V*_nd_ from baboon brain (Horti *et al*, 2014) yielded the binding potentials necessary to estimate in vivo the average values across all binding regions of *B*_max_ (0.67-0.82 nM)) and *K*_i_ (range 170-385) in human brain. While the value of *V*_nd_ in humans currently is unknown, the baboon value of 5.4 ml/cm^3^ estimated here with SEM of 2.7 ml/cm^3^ corresponded to ranges of *B*_max_ estimates of 0.75 (range 0.38-1.83) nM and DMXB-A *K*_i_ estimates of 138-363 nM. These values agree favorably with estimates postmortem of 0.28 nM *B*_max_ for human cerebral cortex with α-[^125^I]bungarotoxin (Davies and Feisullin, 1981) and cortical *B*_max_ values of 0.7 nM in patients with schizophrenia compared to 1.5 nM in healthy control subjects (Marutle et al., 2001), assuming 10% mg protein converted to wet weight.

### Schizophrenia

In the feasibility study of six patients with schizophrenia, we observed abnormalities of α7-nAChR in five of the six subjects. The finding of significant reduction of regional distribution in five of six subjects in two of three regions after outlier exclusion of a single subject agreed with previous findings post-mortem. However, in the present study, the difference between controls and patients is lower than observed post-mortem. This finding could be due to a number of factors, such as tracer differences ([^18^F]-ASEM vs. [^125^I]-BTX), in vivo vs. in vitro tests, and the heterogeneity of the patient population.

## Limitations

### Antipsychotic medication

Neuroreceptor studies of patients with schizophrenia in the drug free or naïve state is increasingly logistically challenging, although such studies have been completed in limited populations (Farde et al., 1986; Wong et al., 1986). In the present limited population, patients with schizophrenia were tested while on stable medication for more than three weeks with antipsychotics that are held to have only negligible effect on binding to α7 nAchR, and to be non-exclusionary in therapeutic studies of partial agonists such as DMXB-A, e.g., clozapine. Several precautions were taken to reduce the possibility of confounding antipsychotic action in the quantification of binding to α7nAchR. First, the majority of the patients were treated with risperidone and haloperidol, which unlike clozapine fail to normalize the P50 abnormality (Adler et al., 1998). Second, a rodent biodistribution study found no significant effects on [^18^F]ASEM binding. Finally, although only one subject was studied off medications, the repeated PET test on-and off-chronic risperidone was suggestive of the conclusion that this commonly used antipsychotic is unlikely to confound the estimates of the *V*_t_ values for _α7_ nAChR.

### Effects of Smoking

All told, all 21 control subjects and 5 of the 6 patients with schizophrenia were non-smokers. We considered comparing smokers with non-smokers but did not recruit a sufficient number of healthy volunteers with clinically verified smoking status. By design therefore we studied only non-smoking control subjects in the current protocol of test-retest and occupancy studies and comparison with the feasibility study of six patients with schizophrenia.

Despite the relatively high prevalence of smoking in patients with schizophrenia, only one of the subjects randomly recruited with schizophrenia smoked (20 cigarettes a day). Most subjects were younger (< 35 years old), which may be a contributing factor, as it has been reported that younger subjects are less likely to smoke^1^. The patient that happened to be a regular smoker also happened to be the oldest of the patients at age 31 (age range of patients 20-31 years, mean 24.9 years). The *V*_t_ values did not differ significantly for the smoking subject compared to the other patients with schizophrenia with low value of *V*_t_.

Furthermore, on the day of PET, all control subjects and patients with schizophrenia had CO values less than 4 PPM, indicating successful abstinence (regardless of their self-reports) and likely longer than 12 hours, reducing the acute effects of nicotine, if any, on the PET measure of the tracer volume of distribution (*V*_t_).

As future direction, we specifically will recruit participants for inclusion into cohorts of smokers and non-smokers of both healthy controls and patients with schizophrenia, to further explore smoking effects. The assignments will be verified by smoking history, urine cotinine levels, CO breath measures, and assessments of extent of nicotine use and dependence (e.g, Fagerstrom, MINI nicotine use, Minnesota Nicotine Scale, and Michigan Nicotine Reinforcement Questionnaire).

With a larger sample, the schizophrenia group is likely to comprise more smokers, particularly among older patients.

### Age and sex differences

In the group of 21 subjects in the age range of 18 to 50 years, males and females were not distributed uniformly, ruling out formal determination of sex differences as functions of normal aging. Also, the small number of five subjects with DMXB-A occupancy nonetheless yielded a reasonable range of DMXB-A human plasma concentrations, presumably due to individual variability in oral absorption and metabolism of the inhibitor.

### Schizophrenia

In this feasiblity study, the small number of patients with schizophrenia is a statistical limitation. In five of the six subjects, estimates of *V*_t_ in two of three regions compared to age-matched control subjects, as well as a formal outlier analysis, suggested that the high values of *V*_t_ in one subject may have had other causes leading to the elevation. Thus expansion of the number of patients with schizophrenia and detailed documentation of the clinical and neuropsychological features the patients will help expand the findings of factors that influence the heterogeneity of α7nAchR findings.

## Conclusions

Here, we confirmed the test-retest reliability of tracer binding in healthy human brains, and we demonstrated for the first-time specificity of competition from the partial agonist DMXB-A in normal healthy volunteers, as previously reported in preclinical imaging studies. The novel first time presentation of in vivo B_max_ measures agreed favorably with published human postmortem values. The effects of antipsychotics in our schizophrenia volunteers may not be a major confound based on our preliminary mice and literature and one subject on and off risperidone. The preliminary findings in patients with schizophrenia suggest the feasibility of imaging this important disorder and generally consistent with previous reports of human brains post-mortem, but clearly larger samples with a spectrum of clinical severity and comparisons with full cognitive and neuropsychological tests are needed.

## Acknowledgements

Robert F Dannals and Radiochemistry staff, Johns Hopkins PET Center staff, David Schretlen, PhD and J Coughlin, MD for scientific discussions and the High Resolution Brain Imaging Section faculty, staff and fellows. Thanks to Gayane Yenokyan, MD, PhD, Director, Biostatistics Consulting Center, Johns Hopkins School of Public Health for guidance and assistance on statistical analyses.

## Footnotes

1 (for an example of this trend, see https://www.hhs.gov/ash/oah/adolescent-development/substance-use/drugs/tobacco/trends/index.html)

